# Expression of genes in the *16p11.2* locus during human fetal cortical neurogenesis

**DOI:** 10.1101/633461

**Authors:** Sarah Morson, Yifei Yang, David J. Price, Thomas Pratt

## Abstract

The 593 kbp *16p11.2* copy number variation (CNV) affects the gene dosage of 29 protein coding genes, with heterozygous *16p11.2* microduplication or microdeletion implicated in about 1% of autism spectrum disorder (ASD) cases. The *16p11.2* CNV is frequently associated with macrocephaly or microcephaly indicating early defects of neurogenesis may contribute to subsequent ASD symptoms, but it is unknown which *16p11.2* transcripts are expressed in progenitors and whose levels are likely, therefore, to influence neurogenesis. Analysis of human fetal gene expression data revealed that of all the *16p11.2* transcripts only two, *ALDOA* and *KIF22*, are significantly enriched in progenitors. To investigate the role of *ALDOA* and *KIF22* in human cerebral cortex development we used immunohistochemical staining to describe their expression in late first and early second trimester human cerebral cortex. KIF22 protein is restricted to proliferating cells with its levels increasing during the cell cycle and peaking at mitosis. ALDOA protein is expressed in all cell types and does not vary with cell-cycle phase. Our expression analysis suggests the hypothesis that the simultaneous changes in KIF22 and ALDOA dosage in cortical progenitors causes defects in neurogenesis that may contribute to ASD in *16p11.2* CNV patients.

---

Large, recurrent Copy Number Variations (CNVs) are implicated in many neuropsychiatric disorders including autism spectrum disorders (ASD), epilepsy, intellectual disability (ID) and schizophrenia (McCarthy et al. 2009; Girirajan and Eichler 2010; Levy et al. 2011; Sanders et al. 2011; Malhotra and Sebat 2012). The *16p11.2* CNV (OMIM 611913) encompasses a 593 kb DNA sequence in the p11.2 region of human chromosome 16 (BP4-BP5). This region harbors 29 protein coding genes and is strongly linked to neurodevelopmental disorders (NDDs) including ASD (Kumar et al. 2008; Bijlsma et al. 2009; Rosenfeld et al. 2010; Shinawi et al. 2010; Zufferey et al. 2012). This *16p11.2* region is flanked by two homologous 147kbp sequences that arose after the evolutionary divergence of humans from other primates, generating a hot-spot for mis-aligned recombination that explains the high frequency of the *16p11.2* CNV in the human population and also the high frequency of *de novo 16p11.2* CNV (Nuttle et al. 2016). In humans, the *16p11.2* microdeletion is associated with transient infant brain overgrowth and focal thickening of the cerebral cortex, while the *16p11.2* microduplication is associated with reduced brain size (microcephaly) (Qureshi et al. 2014; Blackmon et al. 2018). The early manifestation of anatomical phenotype in newborns, along with the onset of ASD symptoms in infancy, suggests crucial roles for *16p11.2* genes during neural development. *16p11.2* is the most prevalent CNV associated with ASD, ~1% incidence, making this CNV particularly intriguing and providing motivation for investigating the role played by *16p11.2* genes in brain development and function (Weiss et al. 2008). Available lines of evidence from *16p11.2* rodent models, *16p11.2* patient derived lymphoblastoid cell lines, and human induced pluripotent stem cells genetically engineered to harbor the *16p11.2* CNV indicate that all *16p11.2* mRNAs’ levels reflect the altered gene dosage of 16p11.2 genes (50% in microdeletion and 150% in microduplication heterozygotes) (Horev et al. 2011; Blumenthal et al. 2014; Pucilowska et al. 2015; Tai et al. 2016). This indicates that multiple *16p11.2* transcript levels are affected by the *16p11.2* CNV and that the pathology of the *16p11.2* CNV could stem from altered dosage of one or more them.

While none of the individual *16p11.2* genes have been identified as sole causative genes for the *16p11.2* phenotype, the use of different models has implicated individual CNV genes in various phenotypes. *MAPK3*, *QPRT, KCTD13*, *ALDO*A, and *KIF22* have been individually associated with a variety of neural phenotypes including cell proliferation, neuronal morphology, axonal projection and spine morphogenesis, altered head size, and behavioral phenotypes. A study in which *16p11.2* genes were systematically knocked down in zebrafish embryos reported neuroanatomical phenotypes for the vast majority of *16p11.2* genes tested, suggesting that the 16p11.2 phenotype is likely to be polygenic (Blaker-Lee et al. 2012; de Anda et al. 2012; Golzio et al. 2012; Pucilowska et al. 2015, 2018).

The cellular mechanisms by which the CNV causes the patient phenotype are poorly understood. One plausible hypothesis is that disrupted neurogenesis causes changes in neuronal output that produces a brain with abnormal cell number or composition and that this contributes to the *16p11.2* pathology. Consistent with this hypothesis, the *16p11.2* deletion mouse model exhibits proliferation defects in cortical progenitors during pre-natal brain development and subsequently develops ASD-like symptoms (Horev et al. 2011; Pucilowska et al. 2015). However, it is unknown which of the proteins produced by *16p11.2* CNV genes are expressed by progenitor cells in the developing human cerebral cortex and are therefore candidates for regulating neurogenesis.

We focused on the potential for the *16p11.2* CNV to affect neurogenesis in developing human cerebral cortex by identifying *16p11.2* genes that are highly expressed in cerebral cortex progenitors and down-regulated as cells become post-mitotic. We analysed previously published human fetal cortex single cell RNA sequencing (scRNA-seq) data (Pollen et al. 2015) to identify candidate genes and characterize their expression in sections of developing human fetal cerebral cortex from the late first and early second trimester.

## Material and Methods

### Human Tissue

Human embryos ranging in age from 12-16 post-conceptual weeks were obtained from the MRC/Wellcome-Trust funded Human Developmental Biology Resource at Newcastle Univeristy (HDBR, http://www.hdbr.org/) with appropriate maternal written consent and approval from the Newcastle and North Tyneside NHS Health Authority Joint Ethics Committee. HDBR is regulated by the UK Human Tissue Authority (HTA; www.hta.gov.uk) and operates in accordance with the relevant HTA Codes of Practice.

For cryosections 12 PCW week brains were fixed in 4%PFA/PBS for 1 week then cryoprotected with 30% sucrose/PBS and then embedded in 50:50 30%sucrose:OCT, flash frozen and sectioned at 12μm using a Leica Cryostat.

The following ages were used for this study: 12 PCW (2 brains), 14 PCW and 16 PCW.

### scRNA-seq Analysis

The publicly available scRNA-seq data set (Pollen et al. 2015) was used to identify candidate genes. Prior to dataset publication the reads were aligned, we normalized the RPKMs as log(x+1). Analysis was performed using R studio. To determine genes with significant changes a Wilcox test by *FindAllMarkers* in Seurat package was used. Monocle2 R package was used to order cells in pseudotime. To identify cell-cycle phase specific transcripts we used function CellCycleScoring from Seurat R package.

### Immunohistochemistry

Immunohistochemistry was carried out on paraffin sections obtained from HDBR. Antigen retrieval consisting of boiling sections in 10mM sodium citrate pH6 for 10 mins was used for all stains. Primary antibodies were diluted in 20% blocking serum in pH7.6 Tris buffered saline (TBS) and sections incubated overnight at 4°C. Primary antibodies used: KID 1/5000 DAB, 1/2000 fluorescent (Invitrogen PA5-29490), KI67 1/800 (Novus Biologicals NBP2-22112), ALDOA 1/100 (Sigma HPA004177).

For colourmetric stains, sections were incubated 1hr at room temperature with biotinylated secondary antibody (1/200) followed by incubation for 1 hour with ABC (Vector Labs) and developed with diaminobenzidine solution (Vector Labs), washed, counterstained with nuclear fast red, dehydrated and then mounted using DPX.

For immunofluorescence sections were incubated with secondary antibodies 1/200 1 hour room temperature, counterstained with 4’,6-diamidino-2-phenylindole dihydrochloride (DAPI;ThermoFisher) and mounted with Vectashield H1400 Hardset Mounting Medium (Vector Labs). Extensive TBS washes were carried out between each step.

### *In Situ* Hybridisation

Cryosections of 12 PCW brain were used for ALDOA and KIF22 *in situ* hybridisation.

PCR primers used to clone *in situ* probes from human cDNA into pGEMTeasy for preparation of DIG labelled RNA were as follows: ALDOA Forward: CTG TCA CTG GGA TCA CCT T, ALDOA Reverse: GTG ATG GAC TTA GCA TTC AC. KIF22 Forward: CGA GAG CGG ATG GTG CTA AT, KIF22 Reverse: GAG ACC CAG GAT GTT TGC CT. *In situ* hybridisation was performed as described previously (Radonjić et al. 2014). Briefly, 12μm cryosections were dried at 37°C for 3 hours then incubated overnight at 70°C in hybridization mix containing 1× salts (200mM NaCl, 10mM Tris HCl (pH 7.5), 1mM Tris Base, 5mM NaH_2_PO_4_2H_2_O, 5mM Na_2_HPO_4_, 0.5M EDTA: Sigma-Aldrich), 50% deionized formamide, 10% dextran sulfate, 1mg/ml rRNA, 1× Denhardt’s, and DIG-labelled RNA probe. Next day sections were washed 3 times at 70°C in wash buffer comprising 1× SSC, 50% formamide, 0.1% Tween-20 and then 3 times at RT in 1× MABT (20mM Maleic acid, 30mM NaCl, 0.5% Tween-20 and pH adjusted to 7.5 with 10mM NaOH). Sections were incubated 1hr RT in 1× MABT blocking solution (20% sheep serum, 2% blocking reagent) and then incubated overnight with anti-DIG antibody 1:1500 in blocking solution at 4°C followed by colour reaction overnight at RT.

### Microscopy and Imaging

DAB and *in situ* hybridisation images were taken using a Leica DMNB microscope with an attached Leica DFC480 Camera. Fluorescence images were obtained with a Leica DM5500B epifluorescence microscope with a DFC360FX camera. Confocal images were obtained using Nikon A1R FILM microscope and analysed in ImageJ.

### Image Analysis and Quantification

For DAB stains and *in situ* hybridisation the images were stitched in ImageJ using the stitching plugin (Preibisch et al. 2009).

For KIF22 analysis of DAB stains rectangular counting boxes (34×88µm) were overlaid across the section. Using ImageJ cell counting plugin cells in each box were counted and denoted KIF22+ (brown) or KIF22-(red). The distinction between the regions (VZ, SVZ, IZ/CP) was determined anatomically by cell density. The count for each box was averaged with other boxes in the region to provide the final value.

For analysis of KIF22/KI67 double staining counting boxes (20×145µm) were overlaid over the VZ and SVZ (determined based on cell density). For determining intensity cells were randomly selected on the DAPI channel, the nucleus outlined and intensity of KIF22 and KI67 recorded. 20 cells were selected per box and the counts from individual boxes combined to give final values.

For subcellular ALDOA analysis counting boxes (20×145µm) were overlaid over the VZ and SVZ. Cells were randomly selected on the DAPI channel, far enough apart to ensure their cytoplasm would not overlap, the Z plane through the center of the cell was used and the nucleus outlined. The KI67 and ALDOA intensity was measured constituting the nuclear value. To obtain ALDOA cytoplasmic intensity the nuclear outline was duplicated and extended 4 pixels allowing a reading of just the cytoplasmic area to be obtained (see Fig.5e). This was performed for 10 cells in each box and the counts from individual boxes combined to give final values.

### Data Analysis and Statistics

Where error bars are shown they are expressed as mean ± SEM. Boxplots show median and upper and lower quartiles. Statistical comparison between two groups was performed with a *t* test. Statistical comparison between more than two groups was performed with ANOCA followed by *post hoc* test. *P<*0.05 was considered statistically significant. Analysis was performed using GraphPad Prism.

## Results

### Analysis of scRNA-seq data identifies *KIF22* and *ALDOA* as progenitor-enriched *16p11.2* transcripts in the developing human fetal cerebral cortex

The *16p11.2* CNV involves microduplication or microdeletion of a 593 kb locus on human chromosome 16 containing 29 protein coding genes (Fig.1a). The aim of the current study is to identify *16p11.2* genes that are potential candidates for being involved in neurogenesis in the developing human cerebral cortex (Fig.1b) and whose altered dosage in *16p11.2* microdeletion or microduplication patients may disrupt neurogenesis and contribute to the CNV phenotype. We reasoned that *16p11.2* genes important for neurogenesis would be highly expressed in proliferating progenitor cells and down-regulated as cells became post-mitotic.

**Fig 1:**
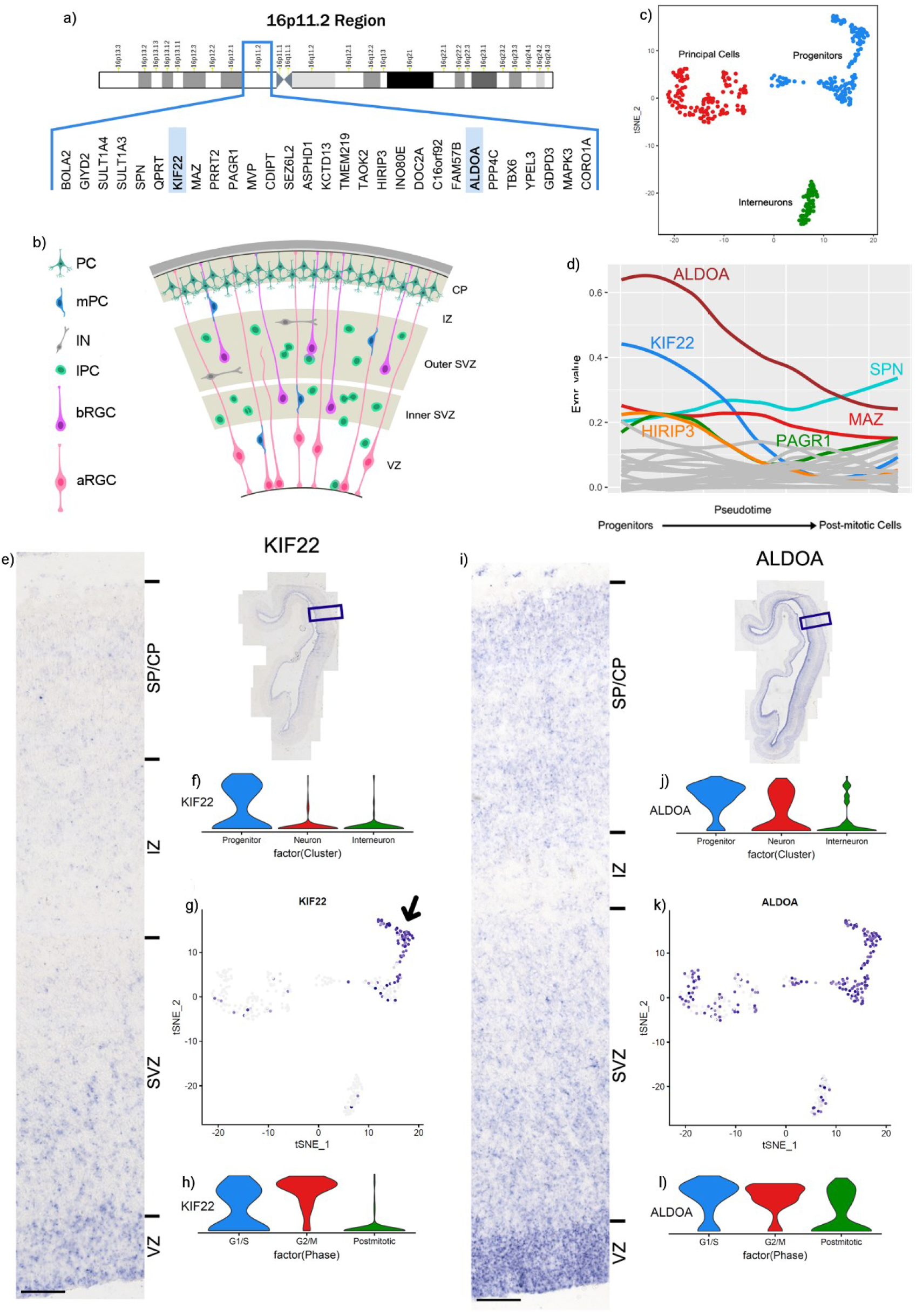
a) 16p11.2 region and genes. b) schematic of human cortical structure during development. c) tSNE clustering of cell types. d) changing mRNA expression levels of 16p11.2 genes as cells move from progenitors to neurons with KIF22 and ALDOA identified as changing significantly. e) *in situ* hybridisation images of KIF22 at 12 PCW. f) Violin plots showing distribution of KIF22 in different cell types. g) KIF22 gradient plot. h) Violin plots showing distribution of KIF22 at different cell cycle stages i) *in situ* hybridisation images of ALDOA at 12 PCW. j) Violin plots showing distribution of ALDOA mRNA levels in different cell types. k) ALDOA gradient plot. l) Violin plots showing distribution of ALDOA mRNA at different cell cycle stages. Scale bars = 100µm.

We took advantage of a published single cell RNA-sequencing (scRNA-seq) data-set acquired from 393 cells of the ventricular zone (VZ) and subventricular zone (SVZ) of gestational week (GW) 16-18 human fetal cerebral cortex (equivalent to post conception week (PCW) 14-16) to perform an unbiased screen to identify *16p11.2* transcripts that matched this expression profile (Pollen et al. 2015). Dimensional reduction of the scRNA-seq data separated the cells into three clusters based on transcriptome similarity (Fig.1c – each dot on the tSNE plot represents an individual cell) that were subsequently identified as the three cardinal cell classes of progenitors (blue), post-mitotic neurons/ principal cells (red) and interneurons (green), by expression of cell-type specific transcripts. We next used the monocle2 R package to order the cells in pseudotime using the normalized expression levels of selected differentially expressed genes (DEGs) as input to order the cells (Trapnell et al. 2014; Qiu et al. 2017) (Fig.1d) moving from the progenitor state (left) to post mitotic state (right) along the X-axis. We plotted the average expression of each *16p11.2* transcript at each pseudotime-point on the Y-axis. We found that two genes, *KIF22* (blue line) and *ALDOA* (brown line), were notable for having high expression in progenitors that declined as cells became post-mitotic. A Wilcox test identified *ALDOA* and *KIF22* as the only *16p11.2* transcripts that were significantly higher in progenitor than neuronal populations (p<<0.05). Although not significantly enriched in progenitors *HIRIP3* (orange line), *MAZ* (red line), *PAGR1* (green line), and *SPN* (turquoise line) transcripts were expressed in progenitors at higher levels than the remaining *16p11.2* transcripts (shown as grey lines), many of which were barely expressed at all.

We focused on *ALDOA* and *KIF22* transcripts and used a combination of *in situ* hybridisation on 12 PCW human cerebral cortex and analysis of the scRNA-seq data to investigate their expression in more detail. *In situ* hybridisation shows that *KIF22* mRNA expression is most prominent in the VZ and SVZ with a few expressing cells in the IZ and SP/CP (Fig.1e). Violin plots of the numbers of cells expressing different levels of *KIF22* mRNA in the different cardinal cell classes show that *KIF22* is expressed predominantly in progenitors (Fig.1f) and mapping the expression level of *KIF22* onto the tSNE plot (Fig.1g) revealed that *KIF22* expression is highest in a subset of the progenitor cluster (arrow in Fig.1g) with a substantial proportion of progenitor cells expressing relatively low levels of *KIF22.* Very few post-mitotic neurons, both interneurons and principal cells, express appreciable levels of *KIF22* (Fig.1f,g). The expression of *KIF22* in a subset of progenitors prompted us to ask whether its expression was related to the cell-cycle phase. We used the expression of cell-cycle phase specific transcripts using function CellCycleScoring from Seurat R package to divide the cells into three classes (Macosko et al. 2015; Tirosh et al. 2016), G1/S, G2/M, and post-mitotic neurons, and compared *KIF22* transcript levels between these three groups using a violin plot (Fig.1h). We found that the majority of cells in G2/M phase expressed higher levels of *KIF22* (red plot), cells in G1/S expressed lower levels (blue plot) while the vast majority of post-mitotic cells expressed low levels of *KIF22* (green plot).

We performed the same analysis for *ALDOA* transcripts. *In situ* hybridisation showed that while *ALDOA* mRNA expression is most prominent in the proliferative VZ and SVZ there are substantial numbers of *ALDOA* expressing cells in the SP/CP (Fig.1i). Violin plots (Fig.1j) show that while a greater proportion of cells expressing the highest levels of *ALDOA* are progenitors (blue plot) there are also a substantial number of principal cells (red plot) expressing similarly high levels of *ALDOA* transcripts although very few interneurons (green plot). Mapping *ALDOA* expression level onto the tSNE plot (Fig.1k) shows cells expressing high levels of ALDOA are evenly distributed throughout the progenitor cluster with appreciable numbers of principal cells expressing high levels of *ALDOA* and a much lower proportion of interneurons. In contrast to *KIF22*, there is no clear difference in the partitioning of *ALDOA* expression level between different phases of the cell-cycle (Fig.1l).

To conclude, of all the 29 *16p11.2* transcripts, only two, *KIF22* and *ALDOA*, are significantly enriched in progenitors compared to post-mitotic cells making them candidates for having specific roles in neurogenesis in the developing human fetal cerebral cortex. Although both are enriched in progenitors, *KIF22* and *ALDOA* transcript expression shows notable differences: *KIF22* transcripts are more restricted to progenitors and their levels vary as the cell-cycle progresses. We next describe the expression of KIF22 and ALDOA protein over a range of developmental stages.

### KIF22 protein is expressed in germinal zones of 12, 14 and 16 PCW cortex

Here we characterize KIF22 protein expression during human corticogenesis. Coronal cortex sections spaced along the rostral-caudal axis were immunostained for KIF22 protein and counterstained with Nuclear Fast Red (NFR) to show cytoarchitecture. KIF22^+^ (brown) and KIF22^−^ (red) cells were counted for each region in the telencephalic wall (VE, VZ, SVZ, IZ and CP) (see methods for details of sampling) and lamination was identified by cell density (Bayer and Altman 2002, 2005). These data are shown for three developmental stages, 12 PCW (Fig 2 a-d), 14 PCW (Fig 2 a’-d’), and 16 PCW (Fig 2 a*-d*). At all stages and rostro-caudal positions KIF22 expressing cells appear most abundant in the VE followed by the VZ and SVZ with the IZ and CP presenting a very low to complete absence of KIF22 (Fig.2 c, c’, and c* with higher magnification of boxed regions from each zone shown in d, d’, and d* respectively, green arrows indicate examples of individual KIF22^+^ cells).

**Fig 2.**
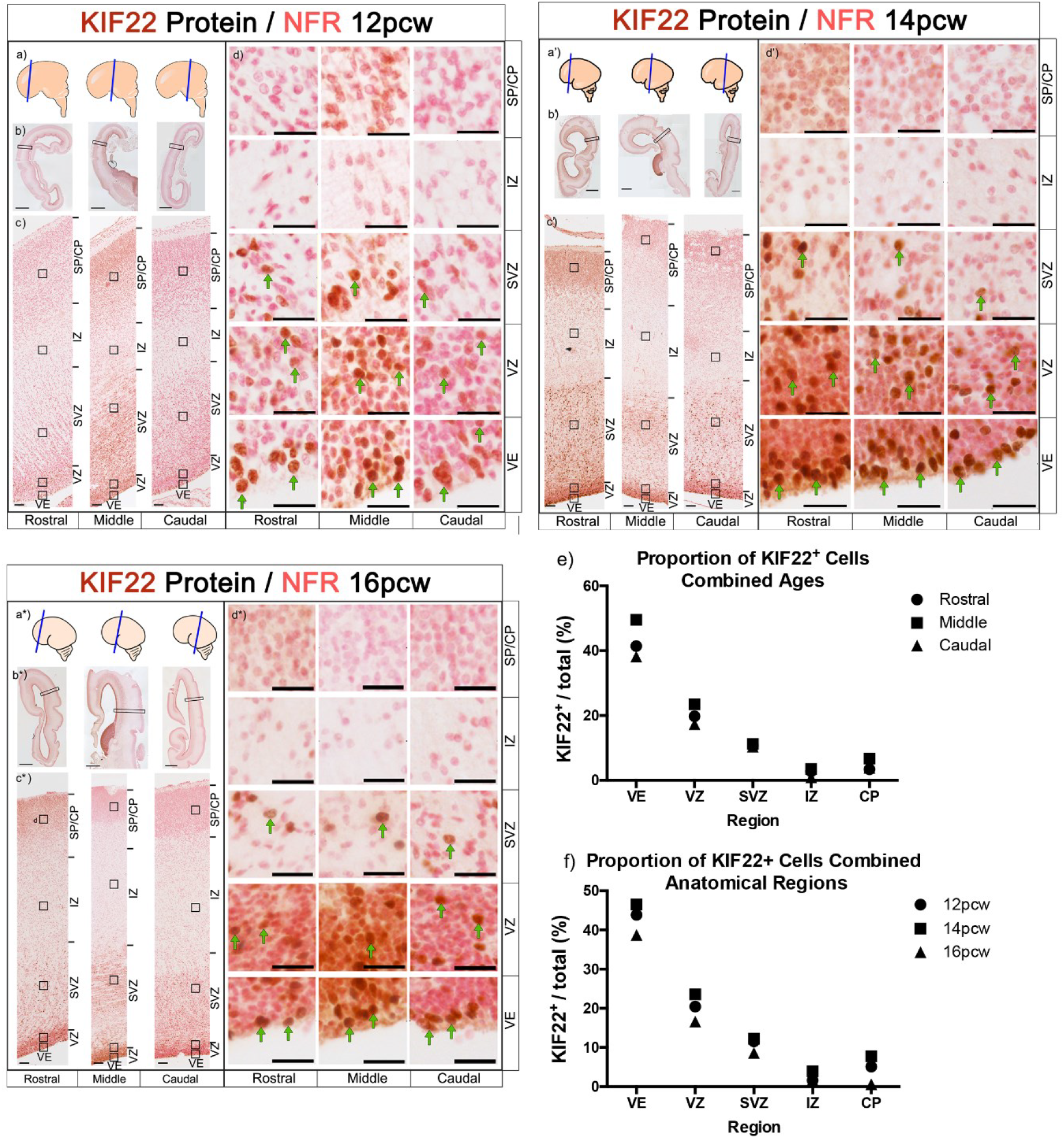
KIF22 protein expression levels in the cerebral cortex at 12, 14 and 16 PCW: a, a’, a*) schematic showing brain regions sectioned. b, b’, b*) images of whole brain section, scale bars =2mm. c, c’, c*) sections spanning the rostral-caudal axis showing KIF22 expression in the telencephalic wall, scale bars = 100µm. d, d’, d*) high magnification images of different cortical zones rostral-caudal. KIF22^+^ cells in brown and examples indicated by green arrows, KIF22^−^ cells in pink, scale bars =25 µm. e) Quantification of KIF22 expressing cells with all three ages combined. f) Quantification of KIF22 expressing cells with rostral, middle, caudal values combined.

We next pooled KIF22^+^ cell count data in two ways to compare between all ages (Fig 2e) and anatomical regions (Fig 2f) and found that the percentage of KIF22^+^ cells in the VE (40-50%) was consistently higher than other regions, followed by the VZ (20-30%) and SVZ (10%), with even fewer cells (<10%) in the IZ, and CP (Fig.2e,f). This result describes KIF22 protein as predominantly restricted to a subset of cells in the germinal zones of the developing cortex at all stages studied.

### KIF22 protein expression is restricted to proliferating cells

KIF22 protein expression is almost exclusively restricted to a subset of cells in the proliferative regions. From the scRNA-seq data, we expect these to be progenitor cells (Fig.1). To identify these KIF22 positive cells we performed double immunofluorescence for KIF22 and KI67 (a protein expressed in all proliferating cells (Scholzen and Gerdes 2000; Miller et al. 2018)). KIF22^+^ cells were predominantly located in the VE, VZ and SVZ (Fig.2), therefore these regions were examined for analysis. Low magnification of KI67/KIF22 staining are shown in 12 PCW (Fig.3a) and 14 PCW (Fig.3b) with higher magnification showing individual cells in Fig.3c. Cell counts for KIF22^+^/KI67^+^ labelled cells show that the majority (80-90%) of KI67^+^ cells also express KIF22 both at 12 (Fig.3d) and 14 PCW (Fig.3e) or across the rostral-caudal axis. Combining the data for anatomical locations and ages revealed significantly more KIF22^+^/KI67^+^ cells than KIF22^+^/KI67^−^ and KIF22^−^/KI67^+^ cells (Fig.3f).

**Fig 3.**
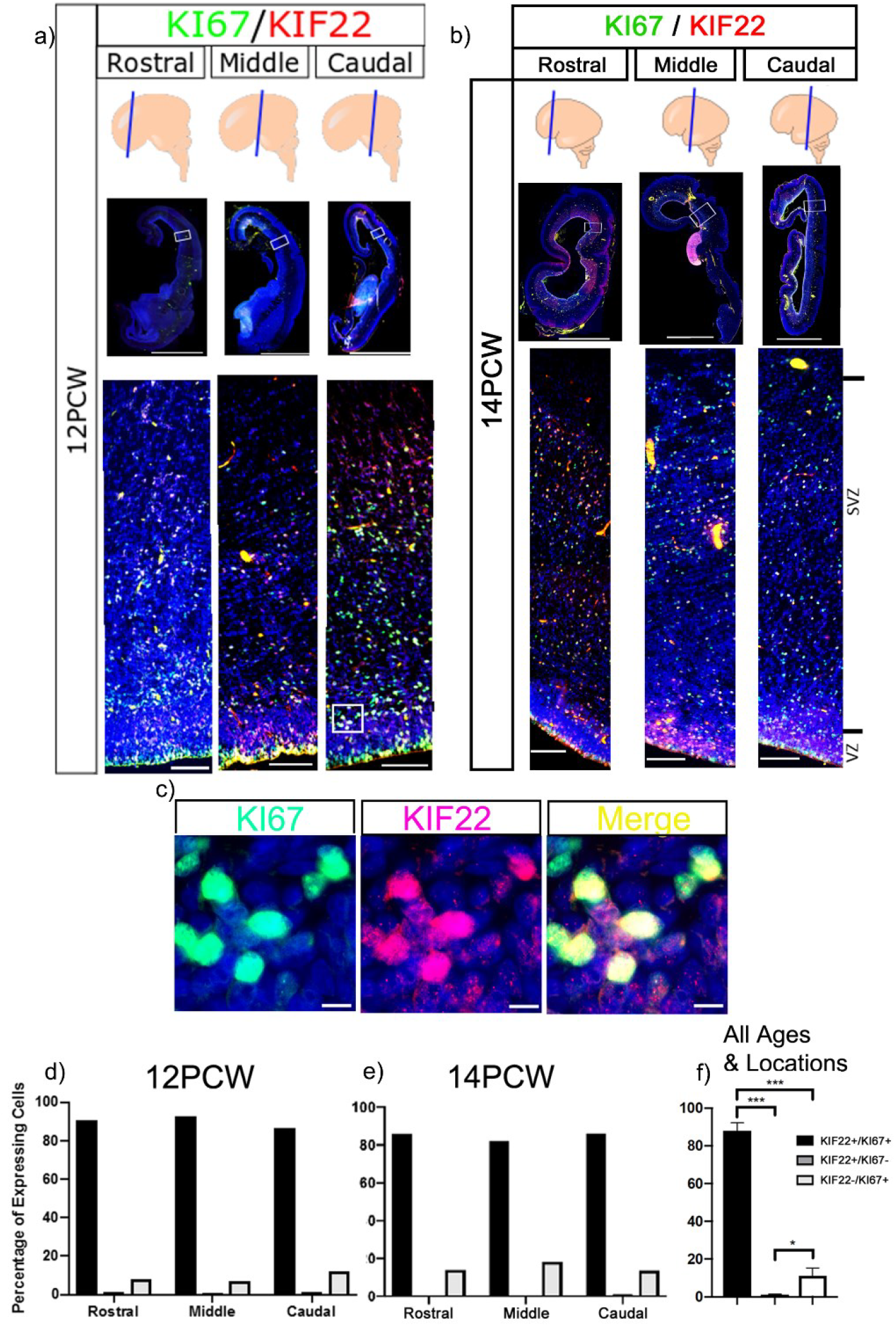
Immunofluorescence of KIF22 and KI67 in the cortex: a) KIF22 and KI67 at 12 PCW, low magnification scale bars = 4mm, high magnification scale bars = 100 µm. b) KIF22 and KI67 at 14 PCW, low magnification scale bars = 4mm, high magnification scale bars = 100 µm. c) high magnification of KI67/KIF22 expressing cells. Scale bars = 10 µm. d) Percentage of cells expressing KIF22, KI67 or both at 12 PCW. e) Percentage of cells expressing KIF22, KI67 or both at 14 PCW. f) Combined data of percentage of cells expressing KIF22, KI67 or both.

Quantification of nuclear KIF22 fluorescence level at the single-cell level shows KIF22 protein is variable, but also significantly higher in KI67^+^ (proliferating) cells than in KI67^−^, across the rostral caudal axis at 12 (Fig.4a-c) and 14 PCW (Fig.4g-i). Combining data from the two ages and locations showed significantly higher KIF22 fluorescence in KI67^+^ cells (Fig.4m). Thus, we confirm KIF22^+^ cells are proliferating and that KI67^+^ cells have higher KIF22 protein levels.

**Fig 4.**
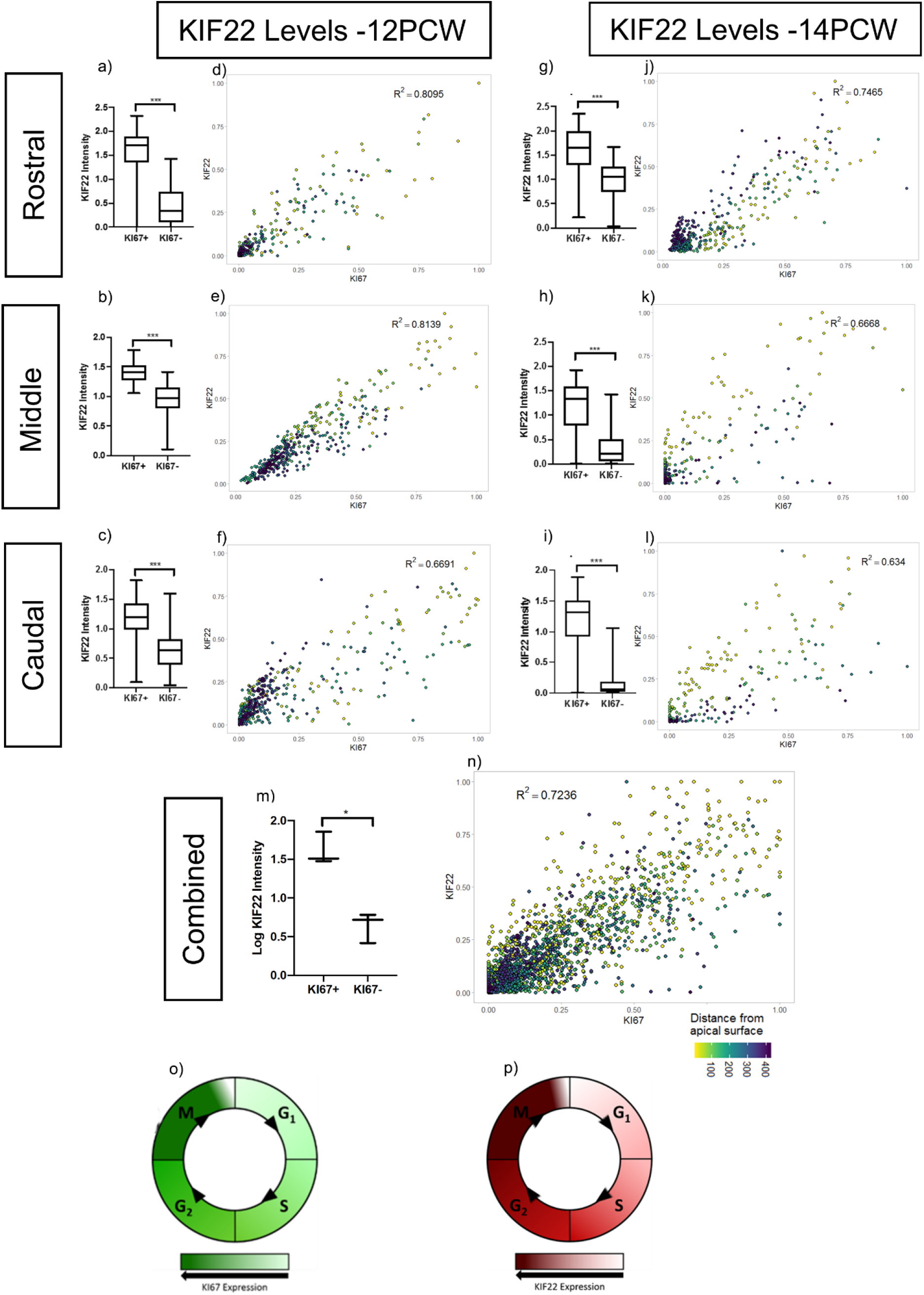
Quantification of KIF22 Levels: a,b,c) 12 PCW quantification of KIF22 fluorescence intensity in KI67^+^/KI67^−^ cells (raw data transformation =+1(log), unpaired t-test with Welch’s correction, p=<0.001). d,e,f) intensity correlations of KIF22 and KI67 nuclear fluorescence intensity at 12 PCW. g, h, i) 14 PCW quantification of KIF22 fluorescence intensity in KI67^+^/KI67^−^ cells (raw data transformation =+1(log), unpaired t-test with Welch’s correction, p=<0.001). j, k, l) intensity correlations of KIF22 and KI67 nuclear fluorescence intensity at 14 PCW. m) quantification of KIF22 fluorescence intensity in KI67^+^/KI67^−^ cells 12 and 14 weeks combined (raw data transformation =(log), paired t-test, p=0.0122). n) intensity correlations of KIF22 and KI67 nuclear fluorescence intensity for rostral-caudal points at 12 and 14 PCW with distance from apical surface indicated by dot colour. o) diagram of KI67 protein levels throughout the cell cycle. p) model based on our results of KIF22 protein levels throughout the cell cycle.

### KIF22 levels vary with cell-cycle phase

From the scRNA-seq analysis, and the variable KIF22 protein levels in KI67^+^ cells, we hypothesised that KIF22 protein levels change throughout the cell cycle. To test this, we quantified nuclear immunofluorescence intensity of KIF22 and KI67 in two 12 PCW brains (see methods for details of sampling procedure). KI67 protein levels vary during the cell cycle: lowest in G1 phase, increasing through S and G2 to peak in mitosis (Fig.4o) (Scholzen and Gerdes 2000; Miller et al. 2018). We found a strong correlation between KIF22 and KI67 intensity (Brain 1 rostral R^2^=0.8095, middle R^2^=0.8139, caudal R^2^= 0.6691. Brain 2 rostral R^2^ = 0.7489, middle R^2^=0.7447, caudal R^2^=0.7763) (Fig.4d-f). To ensure the correlation observed was not a result of nucleus size changing with cell cycle, we confirmed that KIF22 protein levels did not correlate with nuclear size by DAPI staining (Brain 1 rostral KIF22 R^2^=0.104, middle KIF22 R^2^=0.0874, caudal KIF22 R^2^=0.2969. Brain 2 rostral KIF22 R^2^=0.1183, middle KIF22 R^2^=0.0512, caudal KIF22 R^2^=0.2287). A strong positive correlation was also observed at 14 PCW (Brain 1 rostral R^2^=0.7465, middle R^2^=0.6668, caudal R^2^= 0.634, Fig.4j-l). Again, we confirmed that KIF22 protein levels did not correlate with nuclear size (rostral KIF22 R^2^=0.1239, middle KIF22 R^2^=0.0599, caudal KIF22 R^2^=0.0229). This demonstrates that the correlation between KI67 and KIF22 is consistent between ages and rostral-caudal location. Combining all values of KI67/KIF22 nuclear intensity values showed that KIF22 was expressed at significantly higher levels in KI67^+^ cells (Fig.4m) with a strong correlation (R^2^=0.7236) between KIF22 and KI67 levels (Fig.4n). Although KIF22 expressing cells were scattered throughout the VE, VZ and SVZ there was a general trend for the cells expressing the highest levels of KIF22 to be closest to the apical surface (yellow coloured dots on scatterplots Fig.4d-g, g-i,n) with lower expressing cells tending to be further from the apical surface (blue coloured dots on scatterplots Fig.4 d-g, g-I,). During interkinetic nuclear movement radial glial cell nuclei move to the apical surface to perform mitosis so this spatial distribution suggests KIF22 is expressed at high levels by radial glial cells undergoing mitosis at the apical surface of the VZ.

From these data we show that KIF22 protein levels change throughout the cell cycle in positive correlation with KI67: KIF22 is present in G1 and increases through S and G2 phase to peak in mitosis (Fig.4p).

### ALDOA protein is highest in the germinal zones of the cortex

Bioinformatics analysis and *in situ* hybridisation show *ALDOA* mRNA levels decrease as progenitor cells move towards a neuronal fate (Fig.1). Here we used immunofluorescence to characterize ALDOA protein expression across the telencephalic wall at 3 developmental time points; at 12, 14 and 16 PCW, ALDOA immunofluorescence is most intense in the VZ and SVZ before decreasing in the cortical plate (Fig.5 a-c). Double immunofluorescence for KI67 and ALDOA viewed at high magnification shows that ALDOA protein is primarily localized outside DAPI^+^ nuclei in the cytoplasm and that ALDOA is expressed by KI67^+^ progenitor cells and also by cells that do not express KI67 (Fig.5d). The schematic (Fig.5e) illustrates the areas used for quantification of nuclear and whole cell ALDOA fluorescence presented in Fig.6.

**Fig 5.**
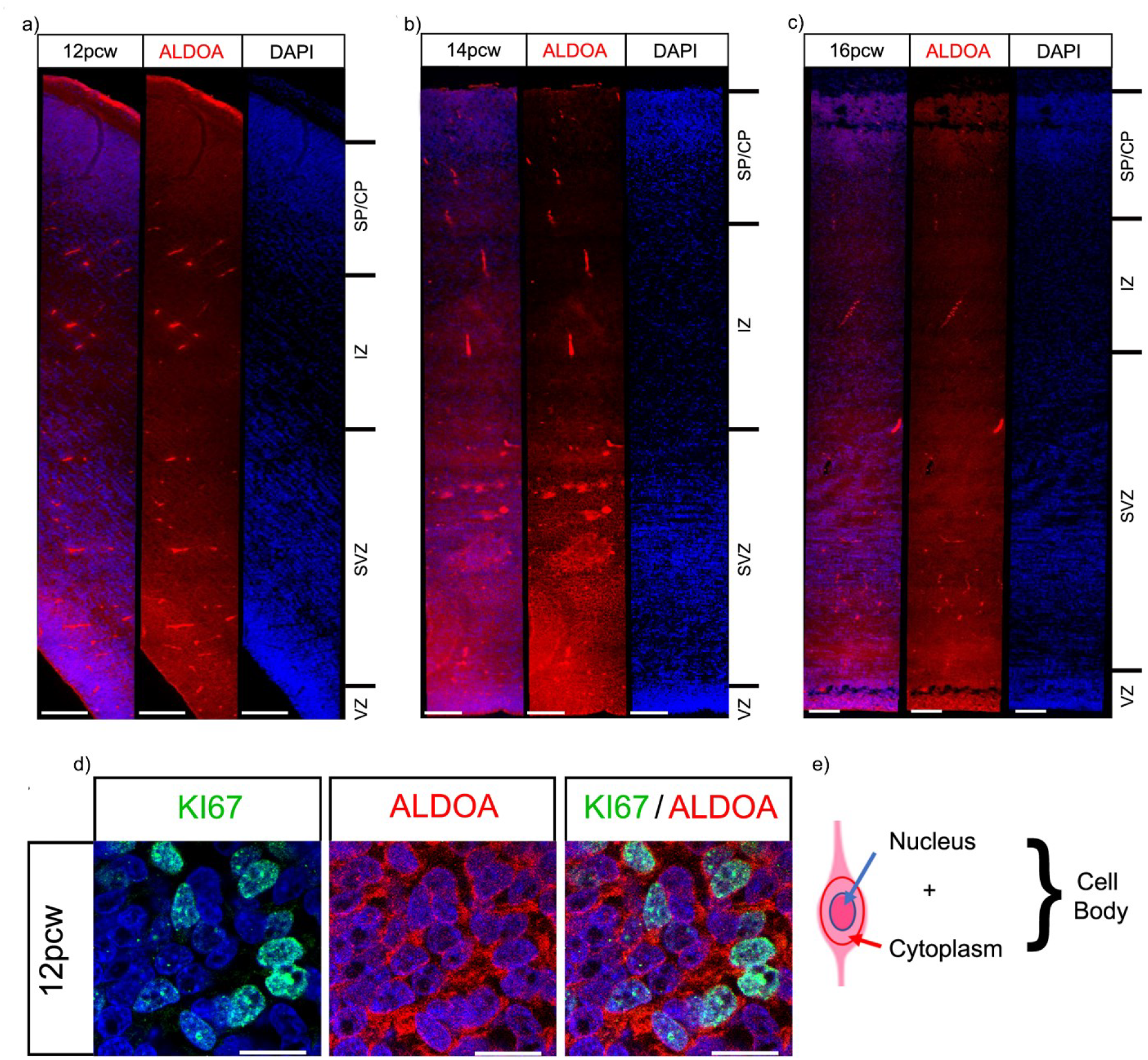
ALDOA Expression in the Cortex: ALDOA protein fluorescence intensity across the telencephalic wall at a) 12, b) 14 and c) 16 PCW. Scale bars = 100 µm. d) High magnification immunofluorescence of KI67 and ALDOA proteins scale bar = 10µm. e) Schematic of quantification method.

**Fig 6.**
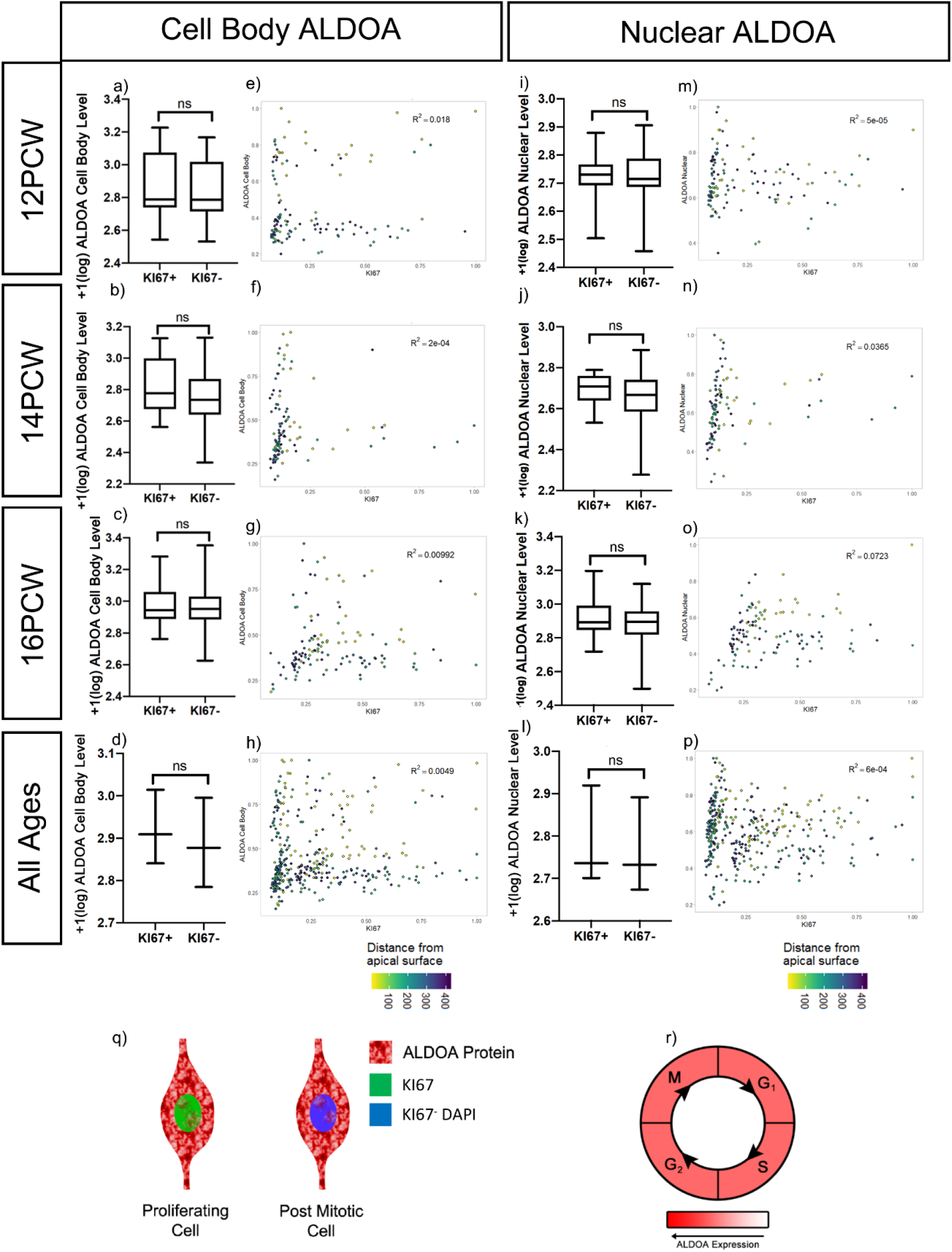
ALDOA Quantification: (a-d) Cell body ALDOA protein fluorescent intensity in KI67^+^ and KI67^−^ cells at a) 12 PCW (raw data transformation =+1(log), bimodal distribution, Mann-Whitney test, p = 0.3702), b) 14 PCW (raw data transformation =+1(log), normal distribution, unpaired t-test with Welch’s correction, p = 0.2032), c) 16 PCW (raw data transformation =+1(log), normal distribution, unpaired t-test with Welch’s correction, p = 0.3523). d) ALDOA cell body protein fluorescent intensity in KI67^+^ and KI67^−^ cells, 12, 14 and 16 PCW individual datasets averaged, (raw data transformation =+1(log)), paired t-test, p = 0.0836. e-h) ALDOA cellular protein intensity levels ls correlated to nuclear KI67 protein intensity at e)12, f)14 and g)16 PCW with distance from ventricular edge indicated. h) ALDOA whole cell protein intensity levels correlated to nuclear KI67 protein intensity pooled 12, 14, 16 PCW. i-l) Nuclear ALDOA protein fluorescent intensity in KI67^+^ and KI67^−^ cells at i) 12 PCW (raw data transformation =+1(log), normal distribution, unpaired t-test with Welch’s correction, p = 0.7543), j) 14 PCW (raw data transformation =+1(log), normal distribution, unpaired t-test with Welch’s correction, p = 0.0694), k) 16 PCW (raw data transformation =+1(log), normal distribution, unpaired t-test with Welch’s correction, p = 0.0772). l) ALDOA nuclear protein fluorescent intensity in KI67^+^ and KI67^−^ cells, 12, 14 and 16 PCW individual datasets averaged, (raw data transformation =+1(log)), paired t-test, p = 0.1330. m-p) ALDOA nuclear protein intensity levels is correlated to nuclear KI67 protein intensity at m)12, n)14 and o)16 PCW with distance from ventricular edge indicated. p) ALDOA nuclear protein intensity levels correlated to nuclear KI67 protein intensity pooled 12, 14, 16 PCW. q) schematic demonstrating ALDOA protein is predominantly in the cytoplasm and lower in the nucleus in both KI67^+^ proliferating cells and KI67^−^ post mitotic cells. r) model showing ALDOA levels do not change with the cell cycle.

### ALDOA protein levels do not correlate with proliferation

Although examination of ALDOA mRNA expression indicated it was enriched in progenitors we were unable to find a significant difference in ALDOA protein levels between KI67^+^ and KI67^−^ cells at 12 (Fig.6a), 14 (Fig.6b) and 16 PCW (Fig.6c) in the human cortex. To look for any fluctuation in ALDOA levels with the cell cycle we quantified immunofluorescence for KI67 and cell body ALDOA (nucleus and adjacent cell body) using the same analysis as that described above for KIF22, and found no correlation or discernible pattern at 12 (Fig.6e) (R^2^= 0.018), 14 (Fig.6f) (R^2^= 2e-4) or 16 PCW (Fig.6g) (R^2^= 0.00992). These data show that in human cortex development, cellular ALDOA protein levels do not correlate with proliferation or fluctuate with cell cycle.

Previous work in different models demonstrated nuclear ALDOA level is greater in proliferating cells (Mamczur et al. 2010, 2013). To see if this was the case in human cortex development, we quantified nuclear ALDOA and KI67 (Fig.5e) but found no significant difference in nuclear ALDOA fluorescence between KI67^+^ and KI67^−^ cells at 12 (Fig.6i), 14 (Fig.6j) or 16 PCW (Fig.6k). We next tested if nuclear ALDOA levels in proliferating cells varied with cell cycle. Analysis of ALDOA and KI67 nuclear intensity established no correlation or pattern at 12 (Fig.6m) (R^2^= 5e-05), 14 (Fig.6n) (R^2^= 0.0365) or 16 PCW (Fig.6o) (R^2^= 0.0723). This shows nuclear ALDOA levels do not increase with proliferation, nor fluctuate with cell cycle. We combined results across the 12, 14 and 16 PCW. There was no significant difference between KI67^+^ and KI67^−^ cells when examining ALDOA protein intensity in the whole cell (Fig.6d) or the nucleus (Fig.6l). Using the pooled data, there was no correlation or discernible pattern when nuclear ALDOA intensity was graphed against nuclear KI67 level for the whole cell (Fig.6h) (R^2^= 0.0049) or the nucleus (Fig.6p) (R^2^= 6e-04).

## Discussion

### *16p11.2* transcript expression during human neurogenesis

The *16p11.2* CNV is a polygenic mutation that causes NDDs and the current study identified a number of the 29 *16p11.2* transcripts expressed in progenitor cells of the cerebral cortex. In addition to *ALDOA* and *KIF22* that are significantly enriched in progenitors, several other transcripts (e.g. *HIRIP3*, *PAGR1*, *MAZ*, and *SPN*) are also expressed in progenitors albeit at lower levels and are not significantly down-regulated as cells become post-mitotic. The simultaneous expression of multiple *16p11.2* genes in cells undergoing neurogenesis suggests that these cells may be particularly vulnerable to simultaneous alteration in their dosage as a consequence of the *16p11.2* microdeletion or microduplication. This lends support to the hypothesis that neurogenesis is disrupted in *16p11.2* CNV patients and that this contributes to subsequent development of NDDs.

### KIF22

KIF22 is a multifunctional protein that can regulate cell proliferation through at least two distinct mechanisms. First, KIF22 is a kinesin-like microtubule-based motor that binds microtubules and chromosomes during mitosis and regulates mitotic spindle microtubule stability and symmetric/asymmetric cell division (Tokai et al. 1996; Tokai-Nishizumi et al. 2005; Sun and Hevner 2014). Second, KIF22 regulates the expression of the cell-cycle regulator CDC25C. During cell division, CDC25C dephosphorylates CDK1, thus activating the CDK1-cyclinB complex while the CDK1-cyclin B complex phosphorylates CDC25C, causing an amplification loop to drive cells to mitosis (Nilsson and Hoffmann 2000). KIF22 directly transcriptionally represses CDC25C and inhibits mitosis; this transcriptional repression of CDC25C is dependent on KIF22 being phosphorylated on Thr463 (Ohsugi et al. 2003; Yu et al. 2014). KIF22 depletion in a tumor cell line accelerates the G2/M transition and slows M/G1 transition (Yu et al. 2014).

Overall it therefore appears that KIF22 can act at several different points in the cell cycle making it difficult to predict how increased or decreased dosage of KIF22 in the *16p11.2* microduplication or microdeletion respectively would impact cell cycle in the specific context of cerebral cortex neural progenitors. Our observation that *KIF22* mRNA and KIF22 protein levels both increase during the cell cycle to achieve highest levels in G2/M phase that drop as cells enter G1 phase implies that KIF22 protein does not persist for long after it is translated and is degraded at the end of M-phase suggesting that both transcriptional and post-transcriptional mechanisms regulate its levels. A clear outcome of our study is that KIF22 levels positively correlate with KI67 in neural progenitors and steadily rise as the cell progresses through G1>S>G2>M phases culminating in the maximum level during M-phase. One possibility is that KIF22 is required to reach a threshold for mitosis to occur, after which its levels must decrease sufficiently to allow mitotic exit. Whether cells undertake proliferative or neurogenic divisions is a process heavily controlled by cell cycle length (Borrell and Calegari 2014). Perturbing *KIF22* gene dosage as a consequence of the *16p11.2* CNV might affect the timing of KIF22 protein reaching this threshold in neural progenitors and therefore affect cell-cycle kinetics and perturb neurogenesis and neuronal output. Our results suggest the hypothesis that KIF22 regulates neurogenesis in the human developing cortex through cell-cycle regulation.

### ALDOA

The process of brain development requires a vast and consistent supply of energy. Glucose is the predominant energy substrate for the fetal brain (Gustafsson 2009), therefore efficient and controlled glycolysis is essential for normal brain development. ALDOA is required for the fourth step of glycolysis, conversion of fructose 1,6-biphospate to dihyroxyacetone phosphate and gluteraldehyde 3-phosphate. The metabolic role of cytoplasmic ALDOA is well established, and ALDOA also has other non-glycolytic “moonlighting” roles such as regulating mitochondrial function and cyctoskeleton stability (Orosz et al. 1988; Pagliaro and Taylor 1992; Kao et al. 1999; Jewett and Sibley 2003; Buscaglia et al. 2006). In addition to its cytoplasmic roles, ALDOA has been observed in the nucleus (Mamczur and Dzugaj 2008; Mamczur et al. 2010, 2013) where it has been suggested to impact cell cycle by positively regulating cyclin D1 expression to mediate G1/S progression (Ritterson Lew and Tolan 2012; Fu et al. 2018). Cell-culture studies show ALDOA sub-cellular localisation depends on the availability of energetic substrates, with addition of glucose driving ALDOA protein to the cytoplasm (Mamczur et al. 2013). Therefore, it is likely the primary role for ALDOA is metabolic when cells require, and have available to them, large amounts of energy. The majority of ALDOA studies have used highly abnormal cancer tissue, or artificial cell culture systems in which glycolytic enzymes have been shown to be increased (Ritterson Lew and Tolan 2012; Mamczur et al. 2013; Fu et al. 2018; Pollen et al. 2019). How these observations of ALDOA in a variety of systems relate to its role in human cerebral cortex development is unclear.

Altering ALDOA dosage in the developing brain will likely impact energy metabolism by altering the flow of metabolites through the glycolytic pathway and impacting subsequent pathways which feed on outputs of glycolysis. Disruption to energy metabolism during development has previously been linked to ASD and ADHD (Rash et al. 2018). The offspring of hyperglycaemic mice presented microcephaly, a phenocopy of the microcephaly observed in *16p11.2* microduplication patients (Rash et al. 2018) and disruptions to energy metabolism may contribute to the microcephaly seen in the offspring of Zika infected mothers (Gilbert-Jaramillo et al. 2019). No homozygous null ALDOA patients have been identified suggesting it is essential for life, but patients with changes to ALDOA levels have been identified; one patient with reduced ALDOA activity presented microcephaly (Kreuder et al. 1996) and another presented intellectual disability (Beutler et al. 1973). Of particular interest is the finding of schizophrenia patients with upregulated cortical ALDOA levels (Beasley et al. 2006) and *16p11.2* microduplication is strongly associated with risk of schizophrenia. This information, coupled with our results that ALDOA is expressed in all cell types, make it clear that any changes to ALDOA dose will perturb energy metabolism at many stages in the brain, impacting its development.

ALDOA is much more abundant in the cytoplasm and we also found no clear relationship between cell body ALDOA levels and cell proliferation status. Nuclear ALDOA has been linked to cell proliferation (Mamczur et al. 2010, 2013; Fu et al. 2018). While we were able to detect nuclear ALDOA we found no clear relationship between ALDOA protein levels and cell proliferation status. While *ALDOA* mRNA levels are higher in proliferating cells compared to non-proliferating cells, quantitative analysis of ALDOA protein failed to identify increased levels in proliferating KI67^+^ cells. This seems unlikely to reflect a technical problem with our quantification method because an identical analysis of KIF22 protein found increased levels of KIF22 protein in KI67^+^ cells (see above). However, our quantification method was restricted to measuring signal in the nucleus and adjacent cytoplasm so may be more appropriate for a nuclear protein (like KIF22) than a protein that fills the whole cell (like ALDOA) and will specifically fail to identify protein localized to the radial fibers of neural progenitors or neurites of post-mitotic cells. Our finding that *ALDOA* mRNA levels do not appear to correlate well with protein levels could therefore have several explanations. One possibility is that *ALDOA* mRNA expression is higher in progenitors and subsequently down-regulated in cells that become post-mitotic but ALDOA protein translated when the cells are progenitors is very stable so persists in post-mitotic cells after *ALDOA* mRNA levels decline. Another possibility is that total ALDOA protein levels are higher in progenitors but the subcellular distribution changes as cells become post-mitotic, for example ALDOA protein in the radial processes of progenitors redistributes towards the nucleus as cells become post mitotic, such that the levels of protein in and around the nucleus remain constant. We cannot distinguish between these possibilities using the fixed post-mortem material available for this study, but it would be possible to test in dissociated cell culture where the whole cell is visible. In any case, it is clear that while ALDOA protein is abundant in progenitor cells of the developing human cerebral cortex and so may play a role in neurogenesis phenotypes, the persistent expression of ALDOA protein as cells become post-mitotic raises the additional possibility that ALDOA also plays roles in differentiated neurons.

## Conclusion

Our study of gene expression in developing human fetal cerebral cortex indicates that further investigation into the normal function of ALDOA and KIF22 and the consequences of altering their dosage on neurogenesis is likely to be a fruitful line of enquiry for understanding the *16p11.2* phenotypes. Further studies are required to unpick the mechanisms involved, but given the nature of the tissue, the scope for studying this *in vivo* is currently limited. However, growth of new model systems in which *16p11.2* gene expression can be manipulated such as human cerebral organoids will provide the opportunity address these questions.

## Acknowledgements

This work was supported by a BBSRC grant (BB/M00693X/1) to TP and an EASTBIO BBSRC funded PhD studentship to SM. The human embryonic and fetal material was provided by the Joint MRC / Wellcome (MR/R006237/1) Human Developmental Biology Resource (www.hdbr.org).

We thank Katherine Howe for her assistance with designing *in situ* probes and undergraduate student Emma Fowler (EF funded by a WR Henderson Scholarship) for her assistance with quantification

